# Resolving Rapid Radiations Within Angiosperm Families Using Anchored Phylogenomics

**DOI:** 10.1101/110296

**Authors:** Étienne Léveillé-Bourret, Julian R. Starr, Bruce A. Ford, Emily Moriarty Lemmon, Alan R. Lemmon

## Abstract

Despite the promise that molecular data would provide a seemingly unlimited source of independent characters, many plant phylogenetic studies are based on only two regions, the plastid genome and nuclear ribosomal DNA (nrDNA). Their popularity can be explained by high copy numbers and universal PCR primers that make their sequences easily amplified and converted into parallel datasets. Unfortunately, their utility is limited by linked loci and limited characters resulting in low confidence in the accuracy of phylogenetic estimates, especially when rapid radiations occur. In another contribution on anchored phylogenomics in angiosperms, we presented flowering plant-specific anchored enrichment probes for hundreds of conserved nuclear genes and demonstrated their use at the level of all angiosperms. In this contribution, we focus on a common problem in phylogenetic reconstructions below the family level: weak or unresolved backbone due to rapid radiations (≤ 10 million years) followed by long divergence, using the Cariceae-Dulichieae-Scirpeae clade (CDS, Cyperaceae) as a test case. By comparing our nuclear matrix of 461 genes to a typical Sanger-sequence dataset consisting of a few plastid genes (matK, ndhF) and an nrDNA marker (ETS), we demonstrate that our nuclear data is fully compatible with the Sanger dataset and resolves short backbone internodes with high support in both concatenated and coalescence-based analyses. In addition, we show that nuclear gene tree incongruence is inversely proportional to phylogenetic information content, indicating that incongruence is mostly due to gene tree estimation error. This suggests that large numbers of conserved nuclear loci could produce more accurate trees than sampling rapidly evolving regions prone to saturation and long-branch attraction. The robust phylogenetic estimates obtained here, and high congruence with previous morphological and molecular analyses, are strong evidence for a complete tribal revision of CDS. The anchored hybrid enrichment probes used in this study should be similarly effective in other flowering plant groups. *[Carex,* coalescent based species tree, flowering plants, low-copy nuclear genes, low-level phylogenetics, universal hybrid enrichment probes]

---

One of the strongest arguments for the use of molecular data in phylogenetic reconstruction was that it provided a seemingly unlimited source of independent characters (Hillis 1987; Hillis and Wiens 2000; Scotland et al. 2003). Although sound in theory, most plant phylogenetic studies are still limited to just two regions, the plastid genome and the nuclear ribosomal DNA (nrDNA; Hughes et al. 2006) region. These regions are widely used because they are easily amplified due to high copy numbers (Álvarez and Wendel 2003) and the availability of universal PCR primers for many of their loci (e.g. White et al. 1990; Taberlet et al. 1991; Baldwin 1992). These regions are also particularly attractive for phylogenetic research because they consist of coding and noncoding loci which evolve at different rates (White et al. 1990; Wicke and Schneeweiss 2015), and they can be used in combination to study processes such as hybridization (Rieseberg et al. 1990; Feliner and Rosselló 2007).

Accessing other sources of molecular characters in plants has not been easy. Although the mitochondrial genome should be a prime source of characters due to high copy numbers, low sequence variation, major structural rearrangements and frequent lateral gene transfer has rendered its sequences impractical for most applications (Palmer and Herbon 1988; Knoop 2004; Bergthorsson et al. 2003; Richardson and Palmer 2006). The plant nuclear genome is equally problematic. Its vast size (63.4-700,000 Mbp; Greilhuber et al. 2006; Pellicer et al. 2010), independent genealogical histories and biparental inheritance are favourable characteristics; however, its generally higher evolutionary rates, low copy numbers and the scarcity of complete model genomes means that designing broadly applicable PCR primers in non-model organisms is rarely successful. This is true even for the most interesting and well-known plant nuclear loci (Hughes et al. 2006) such as the *Waxy* (granule-bound starch synthase) or *LEAFY* genes, and other challenges, such as gene duplication, often necessitate extensive rounds of cloning to differentiate paralogs (e.g., Mason-Gamer et al. 1998; Hoot and Taylor 2001). Even when primers are successfully designed, they often cannot amplify low copy loci from degraded tissue samples, such as herbarium specimens, which means that their usefulness for studies on species-diverse or geographically widespread groups is limited. Consequently, most plant molecular phylogenetic studies consist of only a handful of linked loci from the plastid genome (e.g., *matK, ndhF, trnL-F*) and nrDNA cistron (ITS region, ETS; Hughes et al. 2006) and are rarely comprised of combined analyses of more than five sequenced regions from these two character sources.

Next-generation sequencing (NGS) technologies are directly and indirectly facilitating the exploration of new sources of molecular characters in non-model plants. Although full genome sequencing remains beyond the reach of most systematists, NGS promotes the development of genomic resources for new model organisms, which in turn provides data useful for the design of new Sanger-based markers such as low copy nuclear genes (Blischak et al. 2014; Chamala et al. 2015) or microsatellites (Gardner et al. 2011). In addition, the development of efficient multiplexing and enrichment methods are making NGS increasingly accessible as a method to directly gather data for larger species samples in non-model organisms (Cronn et al. 2012; Lemmon and Lemmon 2013). Low-coverage shotgun sequencing (genome skimming) and organellar genome enrichment permit rapid and efficient sequencing of large phylogenomic matrices from the high-copy regions of genomes (organelles and nrDNA; Straub et al. 2012). However, these approaches are limited by the finite size and generally linked nature of the targeted regions, as they were prior to the invention of NGS. Moreover, they have been unable to completely resolve several important plant radiations (Xi et al. 2012; Barrett et al. 2013, 2014; Ma et al. 2014; Straub et al. 2014). While cost-efficient alternatives, including RADseq (Baird et al. 2008) and transcriptome sequencing (e.g., Wen et al. 2013), can provide data from thousands of unlinked nuclear loci, they both have limitations for phylogenetic analysis. Indeed, RADseq datasets are characterized by short loci of uncertain homology and high amounts of missing data (Rubin et al. 2012; Huang and Knowles 2016), and although it has been used in phylogenetic studies of radiations at least as old as 60 Ma (Gonen et al. 2015; Eaton et al. 2016), the existence of many different protocols and the anonymous nature of RADseq loci (lacking a reference genome) does not facilitate data sharing and reuse across study groups (Harvey et al. 2016). On the other hand, transcriptome sequencing has the potential to be useful at any taxonomic level, but important drawbacks include the complexity of working with RNA (Johnson et al. 2012), especially when living material is not available, and the computational burden of gene assembly and orthology inference in plant genomes where gene families, paralogs, and splice variants are common (Cronn et al. 2012).

A more flexible and promising approach is hybrid enrichment, a method that reduces the bioinformatics and laboratory complexity of transcriptome sequencing by using probes designed from existing genomic or transcriptomic sequences to enrich a fixed set of molecular targets (Lemmon and Lemmon 2013). Low-copy nuclear gene enrichment probes have already been designed to work across vertebrates (Faircloth et al. 2012; Lemmon et al. 2012), and they have been used in the phylogenetic analysis of birds (Prum et al. 2015), snakes (Pyron et al. 2014; Ruane et al. 2015), lizards (Leaché et al. 2014; Brandley et al. 2015; Pyron et al. 2016), frogs (Peloso et al. 2014) and fishes (Eytan et al. 2015) amongst others. In plants, hybrid enrichment probes have been designed for several genera (e.g., de Sousa et al. 2014; Weitemier et al. 2014; Nicholls et al. 2015; Schmickl et al. 2015; Stephens et al. 2015; Heyduk et al. 2016; Johnson et al. 2016), a subfamily of palms (Arecoideae, Arecaceae; Comer et al. 2016), a subfamily of grasses (Chloridoideae, Poaceae; Fisher et al. 2016), and for the sunflower family (Asteraceae; Mandel et al. 2015). Although this taxon-specific approach in plants has been successful, it requires new probes to be designed for every group, and the data generated from such studies has limited potential to be reused because the targeted regions are group-specific. On the contrary, if conserved targets are selected, hybrid enrichment probes can be designed to work on broad taxonomic scales. This method, known as “anchored phylogenomics” (Lemmon and Lemmon 2012), has the potential to provide parallel datasets for a fixed set of loci across large taxonomic groups, like flowering plants. In other words, anchored phylogenomics has the potential to become the modern NGS equivalent of the “universal” PCR primer papers that resulted in an explosion of phylogenetic studies in non-model organisms during the past decades (White et al. 1990; Taberlet et al. 1991; Baldwin, 1992).

The success of hybrid enrichment in several isolated plant groups has motivated us to design a new set of flowering plant-specific probes that can enrich nuclear genes across all flowering plants. In a previous contribution, we identified 517 target loci using 25 angiosperm genomes and we demonstrated their universality and broad utility in flowering plants (Buddenhagen et al., in prep.). This new resource has the potential to greatly simplify and accelerate plant phylogenomic research by reducing the burden of marker choice and probe design, and by promoting the accumulation of parallel data from a standard set of nuclear genes shared by all plant families. Moreover, it would be especially useful if it was able to resolve relationships at both higher and lower taxonomic levels. In fact, the angiosperm probe kit is already being widely adopted through numerous ongoing collaborations to collect data from nearly 90 angiosperm families for more than 2000 samples (to date) at the Center for Anchored Phylogenomics (e.g. Mitchell et al. 2017; http://www.anchoredphylogeny.com). Building upon Buddenhagen et al. (in prep.) where we examine the relationships of major angiosperm lineages, this contribution demonstrates the utility of the method to resolve difficult branches in a rapid radiation of tribes and genera (Cyperaceae, Cariceae-Dulichieae-Scirpeae).

The radiation of *Carex,* the most diverse flowering plant genus of the northern hemisphere (Starr et al. 2015), and its relatives within the Cariceae-Dulichieae-Scirpeae clade (hereafter CDS; Cyperaceae) provides an ideal case to test the utility of universal nuclear gene enrichment probes at medium and shallow phylogenetic depth in plants. With 16 genera and over 2,200 species, this cosmopolitan clade is of considerable evolutionary interest due to its habitat variety (deserts to rain forests), biogeographic patterns (e.g., bipolar, Gondwanan, Amphiatlantic; Croizat 1952), and unique cytology (n = 6 to 56) promoted by agmatoploid chromosomal fusion and fragmentation (Hipp et al. 2009). Previous phylogenies of CDS have identified seven major lineages using the traditional combination of plastid and nrDNA markers (Muasya et al. 2009; Léveillé-Bourret et al. 2014; Léveillé-Bourret et al. 2015). However, like many plant groups, the backbone of the tree remains unresolved possibly due to a relatively old crown age (>40 Ma) and an early radiation that occurred over just 10 million years (Escudero et al. 2013; Spalink et al. 2016). As a result, the most rapidly-evolving plastid genes contain few, if any, characters to support the backbone topology, whereas non-coding plastid and nrDNA regions have diverged so much they cannot be confidently aligned across the whole group. These factors suggest that large numbers of nuclear genes are needed to resolve the backbone phylogeny of CDS, and universal anchored phylogenomics probes could provide the quick and efficient means to obtain them.

The aims of this study were twofold to: 1) test the utility of anchored phylogenomics in closely related genera of flowering plants showing evidence of rapid diversification; and 2) to resolve long-standing taxonomic problems in CDS by estimating a robust phylogeny of the major lineages of the clade. Using the first set of universal probes available for nuclear gene enrichment in flowering plants, we collected data from hundreds of loci in 34 species not included in the initial probe design and representing the full phylogenetic diversity of the CDS clade. To determine whether the results of such an analysis would be compatible with prior analyses, we compared our genomic tree to a phylogeny estimated from a typical plastid plus nrDNA Sanger-derived dataset that is still commonly generated by many researchers today (i.e., a plastid and nrDNA analysis). We discuss the value of anchored phylogenomics for resolving rapid radiations in flowering plants, and the implications of our results on the taxonomy and evolution of the Cariceae-Dulichieae-Scirpeae clade.

## MATERIALS AND METHODS

### Taxon Sampling and DNA extraction

A total of 32 ingroup taxa were included to represent the all major clades of the CDS clade as based on a previous phylogenetic study with extensive taxonomic sampling of the clade (Léveillé-Bourret et al. 2014). This includes *Dulichium arundinaceum* (Dulichieae; comprising ca. 7 spp.), *Khaosokia caricoides* (incertae sedis), both *Calliscirpus* species (Calliscirpus Clade; 2 spp.), *Amphiscirpus nevadensis* (Zameioscirpus Clade; comprising ca. 8 spp.), 5 *Scirpus* and 3 *Eriophorum* species (Scirpus Clade; comprising ca. 48 spp.), 2 *Trichophorum* species (Trichophorum Clade; comprising ca. 18 spp.), and 18 *Carex* species (Cariceae; comprising ca. 2,150 spp.) representing all major Cariceae lineages identified by Starr et al. (2015). Two outgroup taxa (*Eleocharis obtusa* (Willd.) Schult.; Eleocharideae and *Erioscirpus comosus* (Wall.) Palla; Cypereae) were selected from the CDS sister group, the Abildgaardieae-Eleocharideae-Cypereae-Fuireneae clade (Muasya et al. 2009). Two accessions of *Scirpus atrovirens* were included to test the repeatability of the hybrid-enrichment methodology. Leaves collected fresh in the field and dried immediately in silica gel were used in whole genomic DNA extractions using the silica-column based protocol of Alexander et al. (2007) as modified by Starr et al. (2009). However, increased quantities of leaf tissue (80-100 mg instead of 20 mg) and reagents were used to account for the greater mass of DNA required for NGS protocols. We aimed for 1-3 μg of DNA for hybrid enrichment, although some samples with as little as 0.15 μg of DNA worked very well with our methodology. Voucher information is available in Appendix 1.

### Hybrid Enrichment Data Collection

Data were collected following the general methodology of Lemmon et al. (2012) through the Center for Anchored Phylogenomics at Florida State University (http://anchoredphylogeny.com/). After extraction, genomic DNA was sonicated to a fragment size of ~300-800 bp using a Covaris E220 Focused-ultrasonicator with Covaris microTUBES. Subsequently, library preparation and indexing were performed on a Beckman-Coulter Biomek FXp liquid-handling robot following a protocol modified from Meyer and Kircher (2010).

Briefly, sonication is followed by blunt-end repair using T4 DNA polymerase, two different adapters are ligated to both ends of the DNA molecules using T4 DNA ligase, and indexes and full length adapter sequences are added by amplification with 5′-tailed primers. An important modification of this protocol is the addition of a size-selection step after blunt-end repair using SPRI select beads (Beckman-Coulter Inc.; 0.9 × ratio of bead to sample volume). Indexed samples were then pooled at equal molarities (typically 16-18 samples per pool), and then each pool was enriched using the Angiosperm v. 1 kit (Agilent Technologies Custom SureSelect XT kit), which contained probes for 517 flowering plant exons (average: 287 bp, median: 225 bp) as described by Buddenhagen et al. (in prep.). Briefly, the probes were designed by selecting genes that are putatively single copy in *Arabidopsis,* poplar (*Populus*), grape (*Vitis*) and rice (*Oryza*), filtering out exons below the minimum size necessary for enrichment, and then narrowing down on the exons that had ≥55% similarity between *Arabidopsis* and rice. Using these two taxa as reference, orthologous regions from 33 complete flowering plant genomes were identified, and the 517 exons that had an average copy number ≤1.2 per genome were selected for probe design. More details, including the probe sequences, can be found in Buddenhagen et al. (in prep.).After enrichment, 3-4 enrichment reactions were pooled in equal quantities for each sequencing lane and sequenced on paired-end 150-bp Illumina HiSeq 2500 lanes at the Translational Science Laboratory in the College of Medicine at Florida State University.

### Assembly

Reads passing quality filtering were checked for overlap and merged following Rokyta et al. (2012). Reads that could not be merged were treated as unpaired during assembly. Reads were assembled on the flowering plant anchored enrichment references following Buddenhagen et al. (in prep.) and contigs were extended into flanking regions using a *de novo* assembler. Briefly, preliminary matches between each read and the reference sequences were called if 17 bases matched a library of spaced 20-mers derived from the references. Reads were then considered mapped if 55 matches were found over 100 consecutive bases in the reference sequences (all possible gap-free alignments between the read and the reference were considered). The approximate alignment position of mapped reads were estimated using the position of the spaced 20-mer, and all 60-mers existing in the read were stored in a hash table used by the *de novo* assembler. Then, the *de novo* assembler maps additional reads by identifying exact matches between a read and one of the 60-mers in the hash table. Simultaneously using the two levels of assembly described above, the reference sequences were traversed repeatedly until a pass produced no additional mapped reads, enabling extension of assemblies into variable flanking regions. Contigs were estimated from 60-mer clusters. For each locus, a list of all 60-mers found in the mapped reads was compiled, and the 60-mers were clustered if found together in at least two reads. Each cluster of 60-mers was then used to separate the reads into contigs. Relative alignment positions of reads within each contig were then refined in order to increase the agreement across the reads. Up to one gap was also inserted per read if needed to improve the alignment. In the absence of contamination, low coverage or gene duplication, each locus should produce one assembly cluster. Consensus bases were called from assembly clusters as ambiguous base calls (IUPAC ambiguity codes) only when polymorphisms could not be explained as sequencing error (assuming a 0.1 probability of error and alpha equal to 0.05, Buddenhagen et al., in prep.). Called bases were soft-masked (made lowercase) for sites with coverage lower than 5. Assembly contigs derived from less than 10 reads were removed in order to reduce the effects of cross contamination and rare sequencing errors in index reads.

### Orthology, Filtering and Alignment

Orthology was determined for genes with multiple copies following Prum et al. (2015). Briefly, for each locus, the distance between each pair of contig sequences was computed as the proportion of shared 20-mers. The list of 20-mers was constructed from both consecutive bases and spaced bases (every third base). Contig sequences were then clustered with neighbor-joining (NJ) using this alignment-free distance measure, allowing at most one sequence per species in each NJ cluster. This results in multiple clusters, each containing at most one sequence per species. Each cluster is then treated as a probable paralog. Gene copies were efficiently sorted using their variable flanking regions recovered during extension assembly. Clusters containing fewer than 50% of the species were removed from downstream processing. Finally, alignments of the remaining orthologous sequence clusters were performed with MAFFT v. 7.023b (Katoh 2013), with the --genafpair and --maxiterate 1000 flags utilized, and alignments were trimmed/masked using the steps from Prum et al. (2015) and Buddenhagen et al. (in prep.). Briefly, a sliding window of 20 bp was used to mask regions where <10 sites had the most common character present in at least 40% of the sequences. Sites with fewer than 12 unmasked bases were also removed from the alignments. Because of the relatively deep phylogenetic timescale of this study, many variable sites in the regions flanking the conserved exonic core of the probes were masked because they were too variable to align across all taxa.

After the initial automatic alignment in MAFFT, there remained several obviously misaligned regions that were were not removed by the previous filtering step. We initially tried using Gblocks 0.91b (Castresana 2000) to exclude poorly aligned or highly divergent regions, but no parameter combinations could remove some clearly misaligned regions, whereas apparently well-aligned and informative regions were often excluded. This was probably due to the fact that most ambiguous stretches consisted of a few completely misaligned sequences within well-conserved blocks, a situation which is known to confound Gblocks (Castresana 2000). In consequence, all nuclear gene alignments were visually examined and sites containing misaligned bases, diagnosed by long (>3 bp) stretches of disagreements to the consensus sequence in one or a few taxa, were excluded. All the separate alignments were combined in a single concatenated alignment for concatenated analyses or kept separate for analyses based on gene trees.

### Phylogenetic Analyses

#### Parsimony analyses

Heuristic maximum parsimony (MP) searches on the concatenated alignment were performed in PAUP* v4.0 (Swofford 2003) using 1,000 random addition sequence (RAS) replicates, tree-bisection-reconnection branch swapping, holding 5 trees at each step and with the STEEPEST option ON. To prevent undersampling-within-replicate and frequency-within-replicate artefacts, support was assessed with 1000 jackknife 50% replicates using 10 RAS replicates, saving a maximum of 10 trees per RAS, and using the strict-consensus jackknife (GRPFREQ=NO) following the recommendations of Simmons and Freudenstein (2011). Single-locus parsimony analyses and partitioned, hidden and total Bremer support values were calculated in PAUP with the help of ASAP, a perl script provided by Sarkar et al. (2008). Concatenated MP analyses excluding outgroups and/or Cariceae were made to determine whether they could be causing long branch attraction problems affecting ingroup topology.

#### Concatenated maximum likelihood analyses

Concatenated maximum likelihood (ML) analyses were performed in RAxML 8.1.11 (Stamatakis 2014) on the CIPRES Science Gateway v3.3 (Miller et al. 2010). The partitioning scheme was selected among all locus subsets with PartitionFinder v1.1.1 (Lanfear et al. 2012) using the relaxed hierarchical clustering algorithm (Lanfear et al. 2014) based only on subset similarity (--weights 1,0,0,0), with --rclust-percent settings of 1%, 2%, 4% and 10% and using the Bayesian information criterion (BIC), with GTR+G as the only allowed model. The best scoring scheme (BIC = 3,277,453.37809) was found at a --rclust-percent setting of 4% and comprised 19 partitions with 2-127 loci and 395-31,991 distinct alignment patterns. RAxML searches were made with 100 randomized maximum parsimony starting trees and the new rapid hill-climbing algorithm (Stamatakis et al. 2007). Branch support was assessed with 100 (standard) bootstrap replicates (Felsenstein, 1985). Single-locus ML searches were done using the rapid hill-climbing algorithm and 200 rapid bootstrap replicates (option -f a) using a python script to input the parameters to RAxML 8.1.21. Internode and tree certainty values were calculated in RAxML 8.2.4 (Salichos et al. 2014).

#### Species tree analyses

Our phylogenetic problem is characterized by very short backbone branches susceptible to gene tree incongruence (Degnan and Rosenberg 2006). However, fully parametric coalescent-based species tree estimation such as *BEAST is too computationally demanding to be used with hundreds of loci and 34 taxa (Ogilvie et al. 2016). We therefore estimated the species tree with ASTRAL-II v4.10.12 (hereafter ASTRAL), a “summary” species tree method that has been shown to be more accurate and less sensitive to gene tree estimation error than alternatives (e.g. MP-EST) in simulation studies (Mirarab et al. 2014; Chou et al. 2015; Mirarab and Warnow 2015). Trees from the single-locus RAxML searches were used as input in ASTRAL, and branch support was assessed with local posterior probabilities (Sayyari and Mirarab 2016) and with 100 gene and site bootstrap replicates (option -g -r). Internode certainty of the branches of this tree were calculated in RAxML 8.2.4 based on all bipartitions of each quartet, and using the lossless adjustment scheme to correct for incomplete gene trees (Salichos et al. 2014, Kobert et al. 2015).

#### Substitution saturation

Site-specific substitution rates were estimated in IQ-TREE 1.5.0 (Nguyen et al. 2015) by re-optimizing a single GTR+G (16 rate categories) model to the whole concatenated alignment using the ML topology found by RAxML, and using the “-wsr” option. Using these site-specific rates, the 1/3 fastest-evolving sites were extracted and tested for substitution saturation. These sites should be dominated by 3^rd^ codon positions and non-coding bases. Substitution saturation was assessed by plotting raw number of transversions and transitions against GTR distances and noting whether a plateau is attained. Additionally, the statistical test of substitution saturation of Xia et al. (2003) was made in DAMBE (Xia and Lemey 2009; Xia 2015) with 1,000 jackknife replicates on subsets of 4, 8, 16 and 32 taxa.

#### Sources of incongruence

Multiple factors such as lateral gene transfer, gene duplication, hybridization, incomplete lineage sorting and gene tree estimation error cause incongruence between gene trees and their associated species tree. Different types of exploratory data analyses were therefore done to pinpoint the sources of incongruence in our dataset, and especially to determine whether the observed incongruence between gene trees is caused by biological factors (hard incongruence) or is simply due to gene tree estimation error (soft incongruence). Robinson-Foulds distances between ML-estimated gene trees and ASTRAL-estimated species trees were calculated with the ape package (Paradis et al. 2004) in R (R Core Team 2016). We made a linear regression of gene tree to species tree distances against the average gene tree bootstrap support.

In addition, bootstrap support of gene tree branches present or absent in the species tree were compared with the help of histograms. Several reduced concatenated MP analyses were made including only loci conflicting with selected backbone branches (Fig. 1) to determine whether combined analyses would give similarly conflicting results, which would be expected in the case of hard incongruence, or whether combined analysis would negate conflict, which should happen if incongruence was simply due to gene tree estimation error. Scatter plots, histograms and linear regression coefficients were drawn and calculated with R.

**FIGURE 1.**
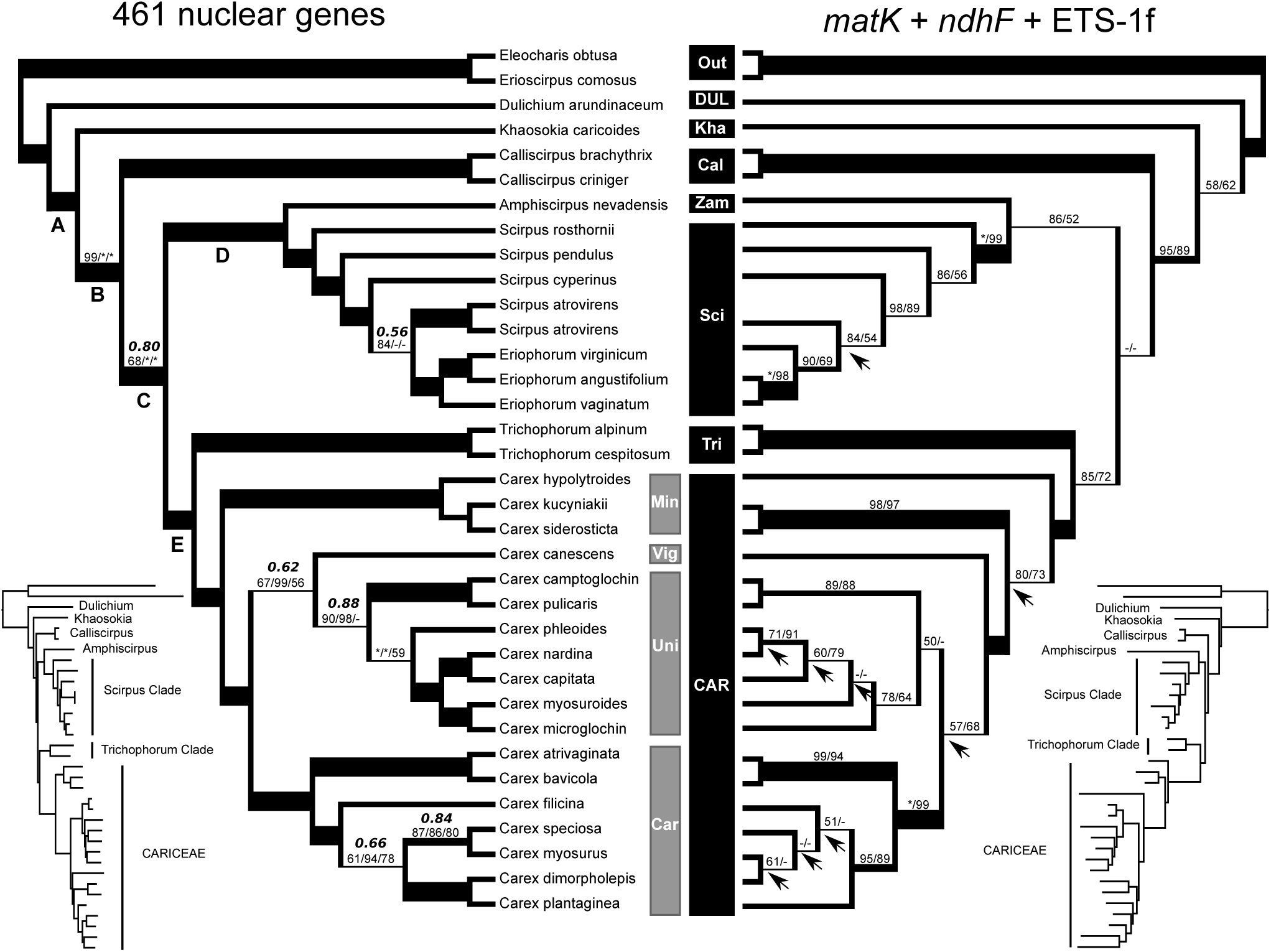
Best phylogenetic hypotheses for the CDS Clade, with the species tree estimated by ASTRAL using 461 NGS nuclear loci on the left, and the ML tree estimated using *matK* + *ndhF* + ETS-1f on the right (arrows indicate topological differences with the ASTRAL species tree). The smaller trees on either side represent relative ML branch lengths for each dataset. Branches without values have 100% support for all measures used. When at least one measure was <100%, the support values are reported above branches as follows: ASTRAL tree (bold italics: local posterior probabily, normal text: ASTRAL multilocus bootstrap/ML bootstrap/MP jackknife) and ML tree (ML bootstrap/MP bootstrap). An asterisk (*) indicates 100% bootstrap support, and a dash (-) indicates less than 50% bootstrap support. Branch width is a function of support in parsimony. Letters under branches refer to clades in Table 1. Legend; Out: outgroups, DUL: Dulichieae, Kha: *Khaosokia,* Cal: Calliscirpus Clade, Zam: Zameioscirpus Clade, Sci: Scirpus Clade, Tri, Trichophorum Clade, CAR: Cariceae, Eri: Eriophorum Clade, Min: Minor Carex Alliance, Vig: *Carex* subg. *Vignea,* Uni: Unispicate Carex Clade, Car: Core Carex Clade. Scirpeae = Cal + Zam + Sci + Tri.

#### Reduced analyses

The influence of the number of loci and analysis method on the reconstructed phylogeny was assessed with reduced analyses. In each analysis, a number of loci were randomly selected (without replacement) and analyzed either by concatenation in RAxML or with ASTRAL. Then, Robinson-Foulds distance between the resulting tree and our best estimate of the species tree (ASTRAL) was calculated. This procedure was repeated 200 times for 5, 10, 20, 50, 100, 200 and 400 loci in ASTRAL, and 50 times for the same number of loci in RAxML. To determine if the information content of the selected loci has an effect on phylogenetic analyses, we ranked loci based on number of informative characters and repeated the ASTRAL reduced analyses with the 33% highest ranking loci and the 33% lowest ranking loci, making 200 replicate analyses with 5, 10, 20, 50 and 100 loci. Boxplots were drawn with R (R Core Team 2016).

### Comparative Sanger Matrix

To compare phylogenetic results obtained with hybrid enrichment, data from the plastid genes *matK* and *ndhF* (Gimour et al. 2013), as well as the nrDNA region ETS-1f, were obtained from Genbank for the same species as those used in phylogenomics analysis (but replacing the outgroup *Eleocharis obtusa* with E. *acicularis* and including only one *Scirpus atrovirens* accession). Five sequences were newly obtained by PCR and Sanger-sequencing following the procotols in Léveillé-Bourret et al. (2015). Genbank accession numbers and voucher information are available in Appendix 2.

The sequences were concatenated by species, aligned with the MAFFT v7.017b (Katoh and Standley 2013) plugin in Geneious 8.1.7 (http://www.geneious.com; Kearse et al. 2012), and the resulting alignments were corrected by hand. Concatenated maximum likelihood (ML) analyses were done in RAxML 8.2.4 (Stamatakis 2014). The partitioning scheme was selected among all codon and locus subsets with PartitionFinder v1.1.1 (Lanfear et al. 2012) using an exhaustive search (Lanfear et al. 2014) and using the Bayesian information criterion (BIC), with GTR+G as the only allowed model. The best scoring scheme (BIC = 31,271.0585328) comprised three partitions: codons 1 and 2 of *matK* and *ndhF,* codon 3 of *matK* and *ndhF,* and ETS-1f. RAxML searches were done using the rapid hill-climbing algorithm and support estimated with 500 rapid bootstrap replicates (option -f a). Support in parsimony was assessed with 500 bootstrap replicates in PAUP* v4.0, using 10 RAS replicates, saving a maximum of 20 trees per RAS, and using the strict-consensus bootstrap (GRPFREQ=NO).

## RESULTS

### Sequence Characteristics

A total of 462 loci were recovered from the Illumina reads, including 456 targets in singlecopy, three targets duplicated into two paralogs each, and only 59 targets not recovered (11.4% of the 517). A single locus possessed no informative variation and was excluded. The average base coverage of loci before trimming was 0-5576 (mean = 234) across terminals, and maximum number of distinct copies per locus (before orthology filtering) ranged from 1 to 4 (mean = 1.4). After alignment and removal of misaligned bases, loci were 219-1,875 bp long (mean = 649), with 1-80% of their length consisting of flanking sequence (mean = 56%), and each with 26-371 parsimony informative characters (mean = 120). Two hundred seventy-nine loci were missing some terminals, but 99% of all loci had at least 29 (83%) terminals. Each terminal had data for 425-462 loci, averaging 457 loci per terminal (98.9% of all loci). The combined dataset was 299,241 bp long after trimming and exclusion of 6,649 misaligned sites. This included 55,417 (18.5%) parsimony informative characters, 4.7% missing, 0.4% ambiguous bases and a GC-content of 40%. Sequence statistics for each locus are found in online Appendix 1 (available from the Dryad Digital Repository: http://dx.doi.org/10.5061/dryad.55h30). The two *Scirpus atrovirens* accessions had 99.3% identical sites and almost identical coverage, with only 5,865 sites (ca. 1% of total aligned length) present in one accession but absent in the other. No evidence of strong substitution saturation was found in the 1/3 fastest-evolving sites. A non-linear trend was visible in the GTR vs transversions and the GTR vs transition plots (online Appendix 2), but the Iss values (0.364-0.424) were significantly smaller (p < 0.001) than critical thresholds (0.603-0.860) for all subset sizes and for symmetrical and asymmetrical topologies, which suggest little saturation (Xia et al. 2003).

### Phylogenetic Results

Concatenated parsimony searches on the phylogenomic matrix found a single shortest tree of 219,733 steps (consistency index = 0.71, retention index = 0.74; online Appendix 3). The best tree found by concatenated ML searches had a log-likelihood of −1,636,930.180799 as calculated by RAxML (online Appendix 4). The best MP and ML trees were almost identical to the ASTRAL species tree (Fig. 1), except for an unsupported sister-relationship between *Carex canescens* (representing the Vignea Clade) and the Core Carex Clade in MP, and a highly supported (100% BS) sister-relationship between *Scirpus cyperinus* and S. *atrovirens* in MP and ML. Relationships between major lineages of the CDS clade, the focus of this study, were identical in all analyses, and their relative position remained stable when outgroups and/or Cariceae were excluded from MP analyses. The MP and ML trees obtained with the comparative Sanger matrix *(matK* + *ndhF* + ETS-1f) were completely congruent with the phylogenomics results, except that most backbone branches had low support in the the Sanger matrix, but very high support in the phylogenomic matrix (Fig. 1).

Phylogenetic analyses position Dulichieae and *Khaosokia* as successive sisters to a highly supported Cariceae + Scirpeae clade. Within this clade, Scirpeae forms four major lineages in three monophyletic groups: a Calliscirpus Clade (*Calliscirpus*) sister to everything else, *Amphiscirpus* (representing the Zameioscirpus Clade) sister to a Scirpus Clade (*Scirpus* + *Eriophorum*), and a Trichophorum Clade (*Trichophorum*) sister to Cariceae. These backbone relationships are highly supported by all analyses except for the position of *Calliscirpus,* which is highly supported in MP and ML, but is supported in only 68% of ASTRAL bootstrap replicates, with the most frequent conflicting BS replicate trees (31%) putting *Calliscirpus* sister to the clade comprising the Zameioscirpus Clade + Scirpus Clade.

### Incongruence Between Gene Trees

Incongruence between ML estimated gene trees was high in the short backbone branches of the phylogeny. This is reflected by small internode certainty values and relatively high numbers of loci with negative partitioned Bremer support (supporting conflicting clades in parsimony) for these short branches identified by letters A to E in Figure 1 (Table 1). However, support and resolution of backbone branches was low in most estimated gene trees, and there was a clear negative relationship between average ML bootstrap (across all branches) of a gene tree and its distance to the ASTRAL species tree (Fig. 2, R^2^ = 0.24, slope = −0.0063, slope p-value < 10^−15^). In addition, the average bootstrap support for branches present in the ML gene trees, but absent in the species tree, was considerably lower than support for branches present in both (Fig. 3). Combined analyses identified extensive emergent support for the backbone branches of the estimated species tree even in loci that are apparently conflicting in single-locus analyses, as shown by the high proportion of hidden Bremer support for backbone branches (Table 1). Likewise, concatenated MP analyses of all loci conflicting with selected backbone branches (identified with letters in Fig. 1) always gave highly supported trees completely congruent with the backbone of our estimated species tree, consistent with soft incongruence due to gene tree estimation error rather than hard incongruence due to biological factors.

**FIGURE 2.**
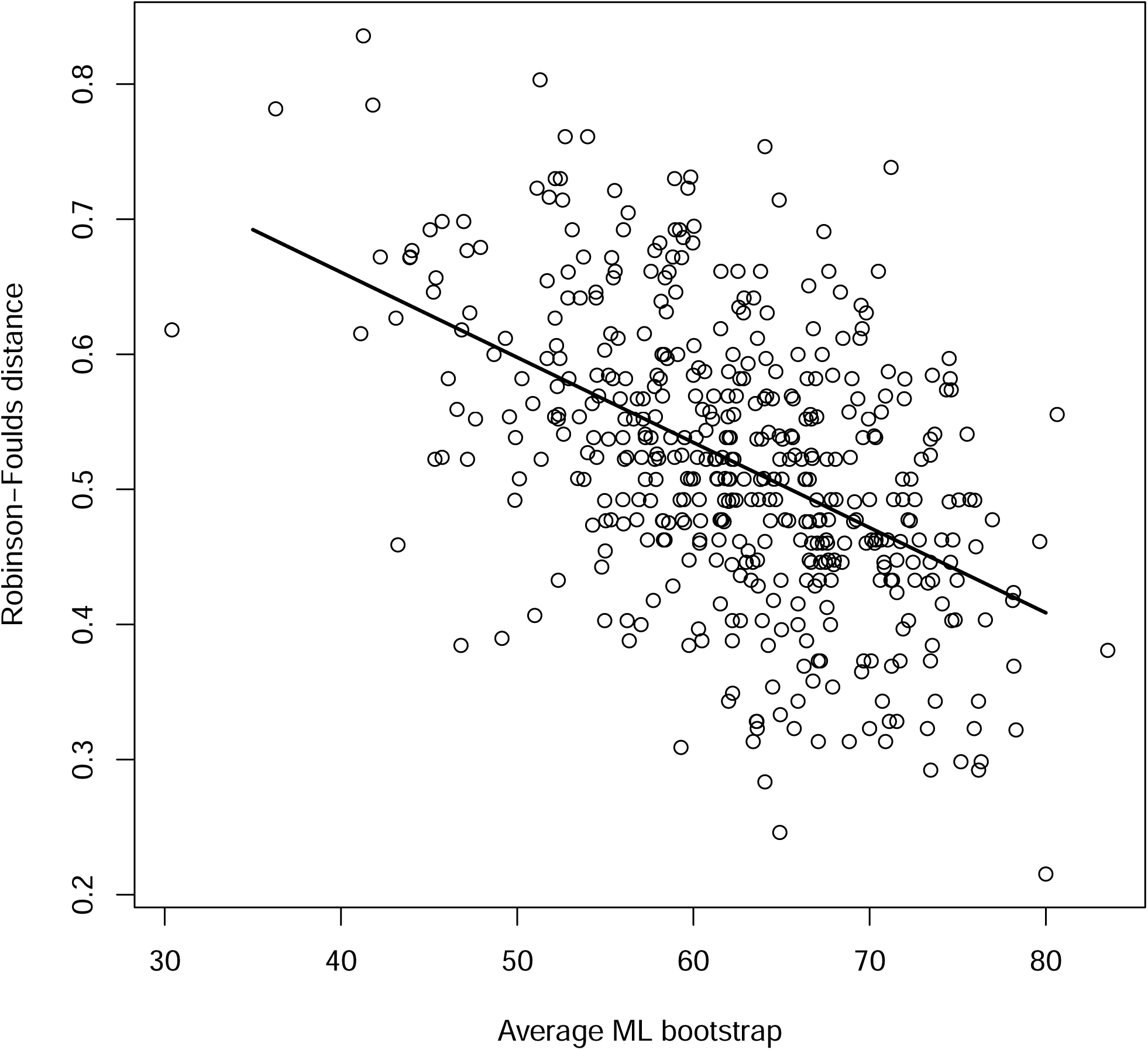
Negative relationship between average ML bootstrap of estimated gene trees and Robinson-Foulds distance between that gene tree and the ASTRAL species tree. Regression line estimated by standard linear regression.

**FIGURE 3.**
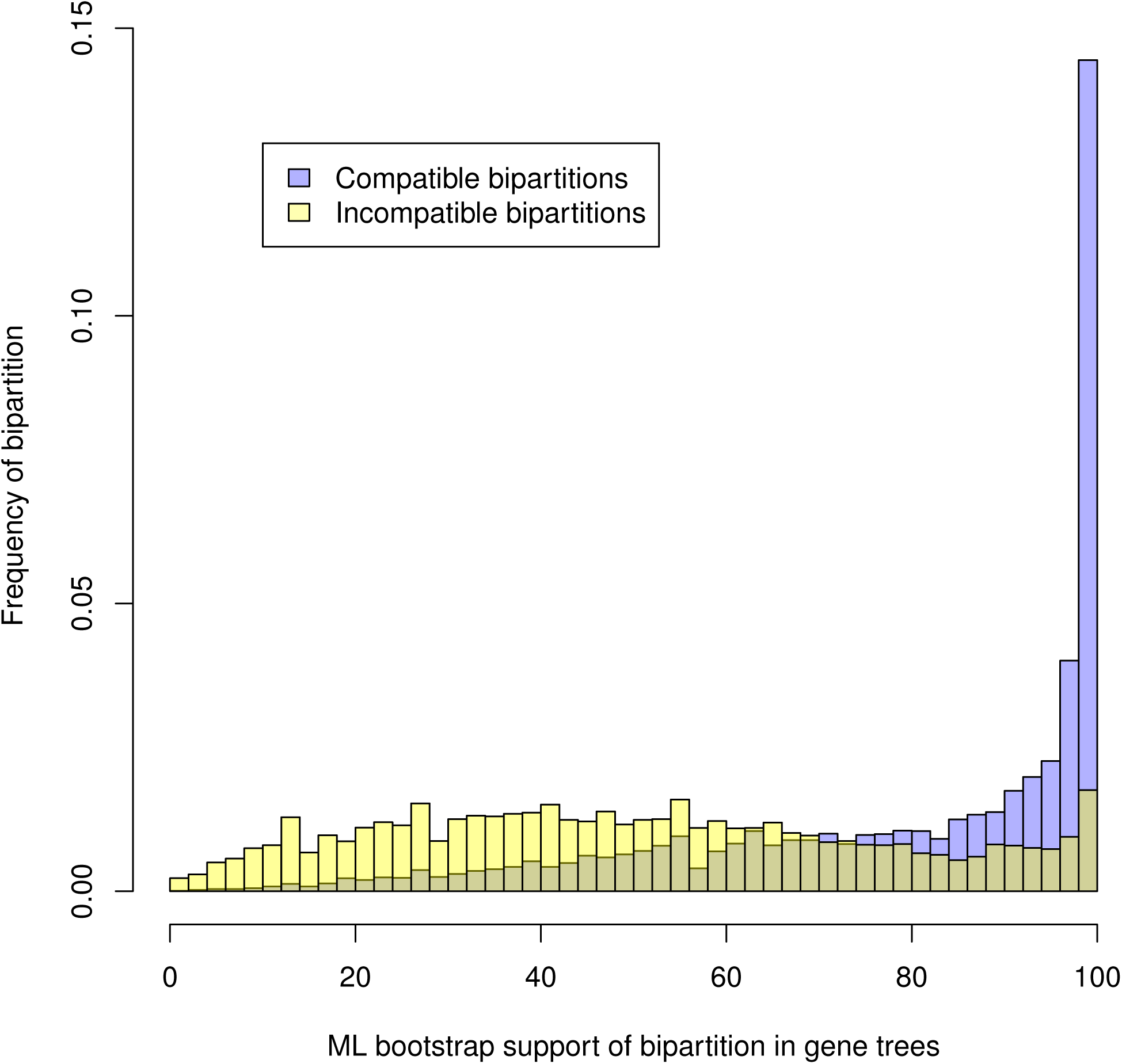
Relative frequency of bipartitions of estimated gene trees as a function of their ML bootstrap support. Bipartitions compatible with the ASTRAL species tree in blue, incompatible in yellow, showing that most gene tree bipartitions are compatible with the species tree beyond about 80% BS.

**Table 1.** Comparison of branch support measures for selected backbone branches (A-E; see Fig. 1). Branch length (expected changes per site in ML), internode certainty (ICA) based on estimated ML gene trees, ASTRAL bootstrap (BS), number of unambiguous synapomorphies, number of loci with positive and negative partitionned Bremer support, total Bremer support and proportion of hidden Bremer support.

| Branch | Branch length | ICA | BS | Unambiguous synapomorphies | Loci positive | Loci negative | Total Bremer | Proportion hidden Bremer |
| --- | --- | --- | --- | --- | --- | --- | --- | --- |
| A | 0.0047 | 0.180 | 100 | 1404 | 296 | 86 | 790 | (43%) |
| B | 0.0023 | 0.066 | 99 | 646 | 209 | 90 | 327 | (66%) |
| C | 0.0014 | 0.006 | 68 | 304 | 195 | 203 | 95 | (43%) |
| D | 0.0058 | 0.380 | 100 | 1082 | 299 | 48 | 921 | (57%) |
| E | 0.0017 | 0.024 | 100 | 364 | 195 | 202 | 137 | (42%) |

### Reduced Analyses

The 33% highest and 33% lowest ranking loci in terms of number of potentially informative characters had an average of 161.2 (sd = 34.4) and 81.6 (sd = 16.7) informative characters, respectively. Results obtained in reduced analyses by using all loci in ASTRAL, RAxML, or with only the highest or lowest ranking loci in ASTRAL, were all broadly similar. Distance between trees estimated in reduced analyses and the best species tree diminished with increasing number of loci per jackknife replicate, with the majority of replicates having less than 10% conflicting bipartitions with 100 loci or more (Fig. 4). With ASTRAL and 200 loci or more, all replicates had a backbone identical to the best species tree, whereas the position of *Calliscirpus* was inconsistent in a minority of replicates with 100 loci. Results were similar when using the highest and lowest ranking loci in ASTRAL. With RAxML, 100 loci were sufficient to get a backbone identical to the species tree in all replicates.

**FIGURE 4.**
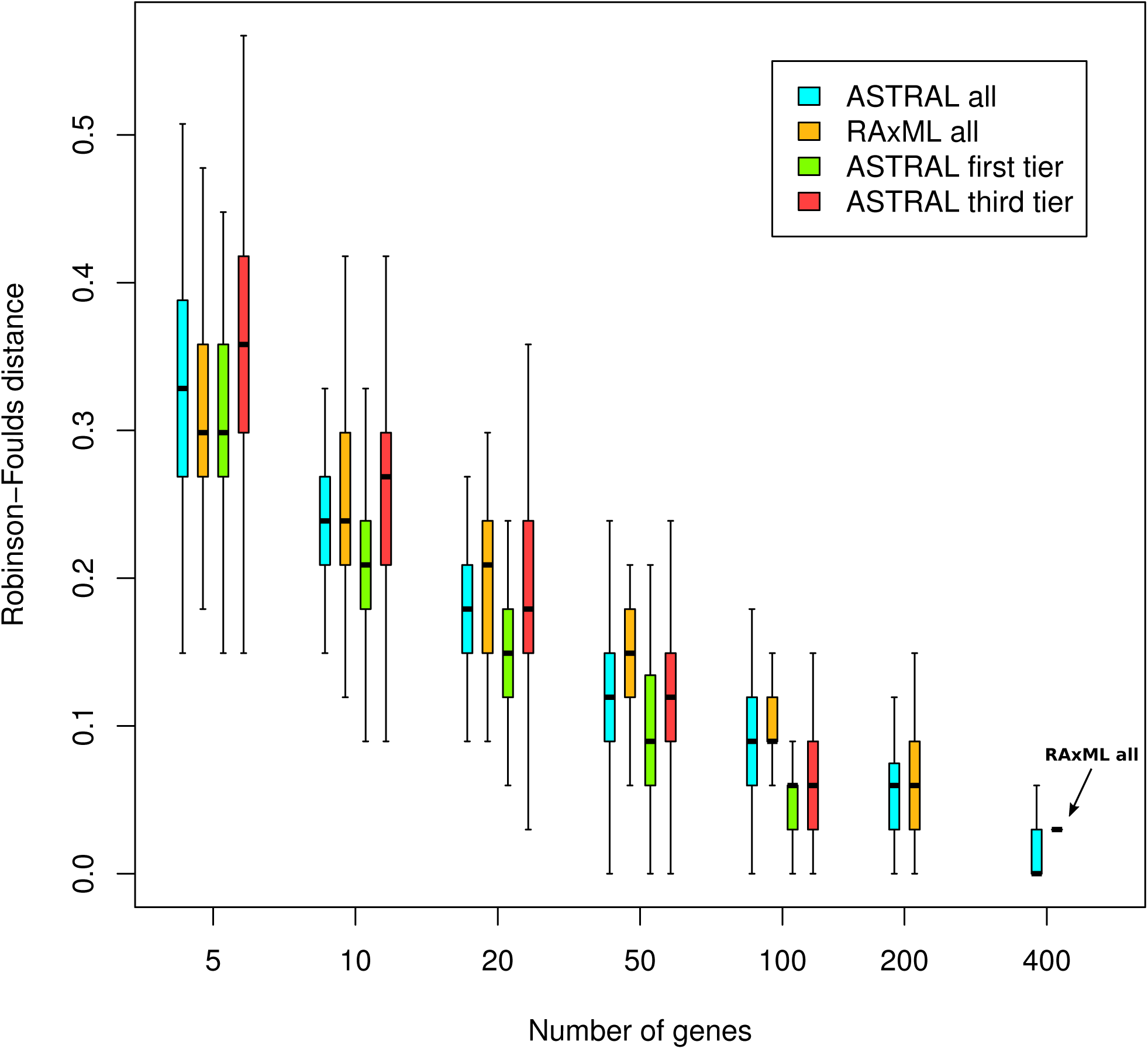
Results of reduced analyses, showing the distribution of Robinson-Foulds distance between reduced analyses tree and the best ASTRAL species tree, as a function of analytic method and number of loci. Note that since all trees were bifurcating, the Robinson-Foulds distance is equivalent to the proportion of conflicting nodes.

## DISCUSSION

### Targeted NGS of Conserved Nuclear Genes in Phylogenetic Inference

Using the first set of universal probes available for nuclear gene enrichment in flowering plants, we were able to collect data from hundreds of loci in 34 taxa representing a typical flowering plant radiation encompassing >40 million years of evolution (Escudero et al. 2013; Spalink et al. 2016). Despite short backbone internodes connected to long branches, typical of ancient rapid radiations (Whitfield and Lockhart 2007), the inferred backbone relationships were well supported in both concatenation and coalescence-based analyses. These results illustrate the great promise of anchored phylogenomics for the resolution of rapid ancient radiations of non-model organisms.

Important amounts of incongruence between estimated gene trees was found in our dataset. The observation that gene tree incongruence was inversely proportional to the amount of phylogenetic information content (as measured by average gene tree ML bootstrap) indicates that at least part of the incongruence must be due to gene tree estimation error. This is corroborated by the lower bootstrap support of gene tree branches absent in the species tree, and the high amount of hidden Bremer support in the shortest branches of the backbone. This also explains why ASTRAL analyses necessitate more loci (ca. 200) than ML analyses (ca. 100) to get consistent results on the backbone relationships of CDS: the estimated gene trees that ASTRAL takes as input are highly affected by estimation error, whereas concatenation presumably amplifies the phylogenetic signal common to all loci, thus reducing the relative influence of noise on the results (Townsend et al. 2012; Bayzid et al. 2015; Warnow 2015; Meiklejohn et al. 2016). The same effect is seen in several simulation studies that have shown higher efficiency of concatenation relative to summary coalescence methods when incomplete lineage sorting is low (e.g. Bayzid and Warnow 2013; Chou et al. 2015; Mirarab et al. 2016). Because the probes used for enrichment were designed to be universal for flowering plants, and since many sites in the variable flanking regions were filtered out because of the phylogenetic depth of the study, the anchored loci tend to be slow-evolving in this study. This resulted in modest numbers of informative characters per locus and low levels of support for individual gene phylogenies. However, it should be noted that the faster evolving flanking regions of the probes could be retained in studies focusing on shallower divergences where additional sequence variation is needed.

It has been argued that small numbers of highly-informative loci are preferable to larger numbers of more slowly-evolving loci when attempting to resolve phylogenies with short branches (Salichos and Rokas 2013). However, the matter is certainly more complex, because more variable loci are often noisier due to multiple substitutions (Townsend et al. 2012) and they have a higher susceptibility to long-branch attraction (Felsenstein 1978; Bergsten 2005). In the case of rapid ancient divergences, difficulties arise because of multiple factors: short backbone branches offer poor phylogenetic signal and increase the probability of deep coalescences, whereas long terminal branches are susceptible to problems of substitution saturation and long-branch attraction (Whitfield and Lockhart 2007). This creates an apparent tradeoff, since fast-evolving loci have a higher probability of containing variation informative for short backbone branches, but are also more susceptible to substitution saturation and long-branch attraction. Indeed, selection of loci should not aim for enormous amounts of variation dominated by noise, but rather for sufficient variation with a high signal/noise ratio and good taxonomic coverage (Philippe et al. 2011; Betancur-R. et al. 2014; Hedtke et al. 2006). For this reason, slowly-evolving, homoplasy-free markers have been suggested to be optimal for the resolution of ancient rapid radiations (Whitfield and Lockhart 2007). Empirical results and simulation studies also indicate that unresolved gene trees are less problematic in a species tree framework than gene trees potentially biased by substitution saturation or long-branch attraction (Chiari et al. 2012; Xi et al. 2015). All these considerations seem to indicate that resolving rapid radiations would profit more from large numbers of conserved loci than from similar numbers of more variable loci. This idea is clearly supported by our success in resolving the polytomy at the base of the CDS clade with hundreds of conserved nuclear genes, despite low levels of support for individual gene trees. Our reduced analyses likewise indicate that using loci with higher or lower numbers of informative characters has almost no effect on our results, whereas the number of loci had a very significant effect on precision of species tree and concatenation analyses. This suggests that future phylogenomic studies based on conserved nuclear loci could profit more from large numbers of loci than from sampling more characters per locus, despite the likelihood that longer loci could decrease gene tree estimation error. However, Meiklejohn et al. (2016) found that species tree estimation methods gave inconsistent results when using gene trees with very low signal (<25 informative characters), whereas more informative loci gave more consistent results. The lack of relationship we found between gene tree information content and species tree accuracy could be explained by the fact that all our loci contained at least 26 informative characters, which suggests that there could exist a threshold below which methods based on estimated gene trees might loose their accuracy. This subject should be explored further to determine whether such a threshold exists.

Our test case suggests that the efficiency of anchored phylogenomics to enrich specific and universal flowering plant loci is highly promising, and the method could thus simplify next-generation sequencing data collection and sharing across diverse flowering plant groups. Of the 517 anchored phylogenomics loci targeted in this study, approximately 90% produced useable data. In addition, different accessions of the same species provided almost identical sequence and coverage, suggesting that the methodology is reproducible and will provide data that can be reused and combined in future phylogenetic studies. Other next-generation sequencing approaches in plants have up to now focused on target-enrichment of lineage-specific nuclear loci (de Sousa et al. 2014; Nicholls et al. 2015; Stephens et al. 2015; Heyduk et al. 2016) or on the anonymous and very short markers provided by RADseq (e.g. Eaton and Ree 2013; Escudero et al. 2014;

Hipp et al. 2014; Gonzalez 2014; Massatti et al. 2016). Lineage-specific target-enrichment approaches have the advantage of being tailored for the group of interest, and are thus expected to perform better on average. However, this is counter-balanced by the additional cost and time needed to design new probes for every taxonomic group, and the limitations that lineage-specific markers impose on data sharing and reuse across taxonomic groups. RADseq, on the other hand, enables rapid and cost-efficient production of tens to hundreds of thousands of loci in large numbers of individuals without the need for genomic references. The short length of RADseq loci (mostly limited by read-length) makes determination of homology difficult, for instance creating a tradeoff between the number of putative loci retained for analysis and the proportion of loci which are truely orthologous (Rubin et al. 2012; Harvey et al. 2016). High levels of missing data due to uneven coverage, mutation-induced locus-dropout or other causes (ca. 30-80% in published analyses; Mastretta-Yanes et al. 2015; Eaton et al. 2016) creates a similar tradeoff where excluding loci with missing data also signicantly reduces the total number of informative characters in the dataset. Despite this, several studies have now demonstrated that radiations at least as old as 60 Ma can be successfully resolved using lax similarity cutoffs during assembly and inclusion of all loci with at least 4 terminals, which suggests that paralogy and missing data may not be problematic for RADseq in most applications (Gonen et al. 2015; Eaton et al. 2016; Huang and Knowles 2016). One advantage of RADseq compared to Anchored Phylogenomics is the ability to tailor the number of loci to the phylogenetic question and ressources available, while there is a hard limit to the number of loci in hybridization-based approaches (517 loci in our case). On the other hand, data sharing and reuse remains an issue with RADseq because of the use of different enzymes and library preparation methods, the difficulty of assessing orthology at deeper evolutionary timescales and the anonymity of RADseq loci when lacking reference genomes (Ree and Hipp 2015; Harvey et al. 2016). Compared to both lineage-specific approaches and RADseq, the universality and easy comparability of anchored phylogenomics results in significant savings in cost and time whilst simplifying data analysis and sharing.

### Phylogenetic and Taxonomic Implications

After nearly two decades of molecular work on the higher-level phylogeny of the sedge family (Cyperaceae; starting with Muasya et al. 1998), one of the most enduring problems has been the placement of tribe Cariceae and the identification of its sister group. This is true despite considerable interest in the evolution, biogeography and ecology of the tribe (more than 140 articles per year since 2010 according to Web of Science), which is derived in part from its exceptional diversity (ca. 2,000 species), global distribution, and peculiar cytology (agmatoploidy, n = 6 to 56, holocentric chromosomes). Lack of knowledge of Cariceae’s sister group has important implications, since the accuracy of any morphological, ecological or geographical character reconstruction is affected by outgroup choice, outgroup relationships, and its effects on ingroup topology (Lyons-Weiler et al. 1998; Wheeler 1990; Graham et al. 2002; Wilberg 2015).

Continued work towards inclusion of more informative molecular regions or increasing taxonomic sampling has resulted in good support for seven major lineages within a clade consisting of tribes Cariceae, Dulichieae and Scirpeae (CDS), but relationships between these major lineages and the identity of Cariceae’s sister group has remained elusive (Léveillé-Bourret et al. 2014). Previous studies have placed Cariceae sister to a monophyletic Scirpeae (Muasya et al. 2009) or nested within a paraphyletic Scirpeae and sister to either a Trichophorum Clade (Léveillé-Bourret et al. 2014), the genus *Calliscirpus* (Gilmour et al. 2013) or a clade consisting of the Scirpus Clade + Zameioscirpus Clade (Jung and Choi 2012). The only consistency has been the poor support for all backbone relationships, a consequence of a rapid radiation (ca. 10 My) followed by long divergence (30-40 My) between major CDS lineages (Escudero et al. 2013; Spalink et al. 2016). Our highly supported results, based on data from hundreds of nuclear genes encompassing hundreds of thousands of base pairs, identify the Trichophorum Clade as sister to Cariceae and are in complete agreement with the results of the most inclusive plastid phylogeny of CDS (Léveillé-Bourret et al. 2014). Such a high congruence between phylogenetic estimates based on the nuclear and plastid genomes gives us confidence in the robustness of the results and confirms the usefulness of targeted-enrichment of conserved nuclear genes for phylogenetic analysis at the tribal level and above in sedges (Cyperaceae). Moreover, since the enrichment probes we used are universal and are flanked by regions of variable evolutionary rates, they could be equally effective for low-level phylogenetic investigation of flowering plants in general.

Our analyses strongly support the paraphyly of tribe Scirpeae, a long-expected result given the likely plesiomorphic nature of its defining characteristics (Goetghebeur 1998). Moreover, the isolated phylogenetic position of *Khaosokia* definitely excludes it from any currently recognized Cyperaceae tribe. These phylogenetic results are congruent with previously identified morphological and embryological variation (Léveillé-Bourret et al. 2014), but a lack of support in previous phylogenetic analyses has prevented taxonomic changes from being made. The robust phylogenetic estimates obtained in this study now provide a solid foundation for a complete revision of the tribal taxonomy of the CDS clade. It is clear that preservation of the highly distinctive Cariceae within a natural and inclusive tribal classification will necessitate the naming of at least three new tribes. Such strongly supported results would probably never have been achieved without genome-scale phylogenetic analyses, which clearly demonstrates the importance of new data acquisition and analysis methodologies in the progress of systematics and taxonomy.

## ACKNOWLEDGMENTS

The authors thank the curators of the following herbaria for loaning material and permission to sample specimens for DNA analysis: Jennifer Doubt (CAN), Dr. Paul Catling (DAO), Dr. Paul E. Berry (MICH) and Dr. Luc Brouillet (MT). We also thank Marie-Ève Garon-Labrecque (DAO), Kellina Higgins (MT), Jacques Cayouette (DAO) and Alexandre Bergeron (MT) for having collected or helped with the collection of canadian material. Nguyễn Thi Kim Thanh (HNU), Vũ Anh Tài (HNU) and Jack Regalado (MO) provided field assistance in Vietnam. Michelle Kortyna, Alyssa Bigelow and Sean Holland help with the lab work. Kirby Birch helped with the bioinformatics. Erika Edwards, Mark Fishbein and five anonymous reviewers made helpful comments on the manuscript. This research was conducted as part of the requirements for a Ph.D. at the University of Ottawa (UofO), with support from an Alexander-Graham-Bell Research Scholarship from the Natural Sciences and Engineering Research Council of Canada (NSERC) and an Excellence Scholarship from the UofO to ÉLB. This research was also funded by a NSERC Discovery Grant to JRS for laboratory and field work, and National Geographic Society Research and Exploration Grants (#9035-11, #9441-14) to BAF (Principal Investigator) for field-work in Vietnam. Additional funding was provided by the National Science Foundation through awards to ARL and EML, (NSF IIP-1313554), and to EML (NSF DEB-1120516).

## Appendix 1 Samples used for anchored phylogenomics

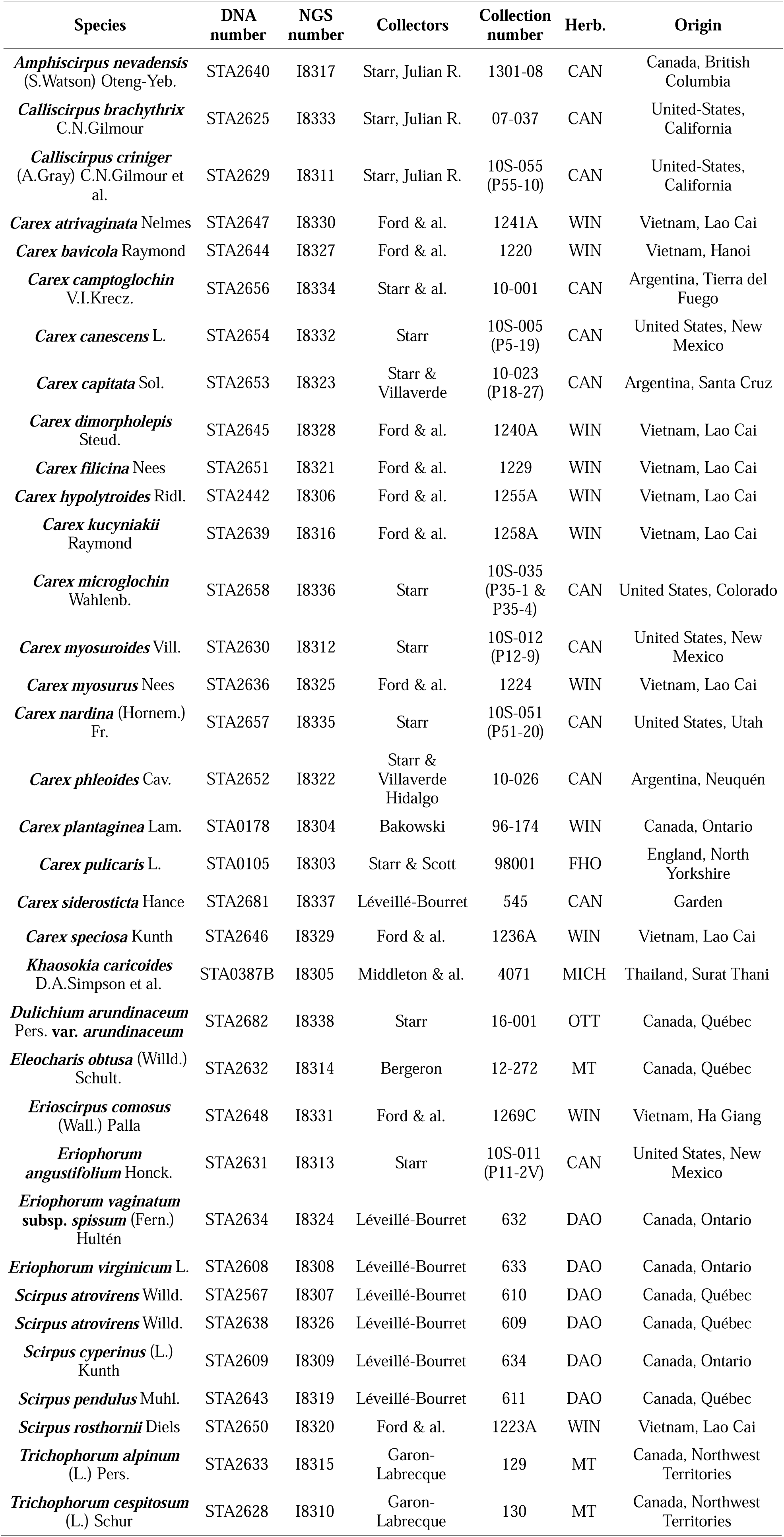

## Appendix 2 Samples and Genbank accession numbers used in the comparative Sanger-based analyses.

APPENDIX 2. Samples and Genbank accession numbers used in the comparative Sanger-based analyses.
| Species | DNA number | Collectors | Collection number | Herb. | Origin | *matK* | *ndhF* | ETS-1f |
| --- | --- | --- | --- | --- | --- | --- | --- | --- |
| *Amphiscirpus nevadensis* (S.Watson) Oteng-Yeb. | STA2141 | Hudson | 5177 | CAN | Canada, Saskatchewan | JX065075 | JX074631 | KP705256 |
| *Calliscirpus brachythrix* C.N.Gilmour | STA2625 | Ahart & Oswald | 5099 | CHS | United States, California | JX065078 | JX074634 | JX065112 |
| *Calliscirpus criniger* (A.Gray) C.N.Gilmour et al. | STA2629 | Chambers | 2973 | DAO | United States | JX074655 | KJ513488 | JX065099 |
| *Carex atrivaginata* Nelmes | STA2419 | Ford & al. | 1230A | WIN | Vietnam, Lao Cai | newly submitted | newly submitted | newly submitted |
| *Carex bavicola* Raymond | STA2389 | Ford & al. | 1215B | WIN | Vietnam, Hanoi | KP273672 | KP273726 | KP273600 |
| *Carex camptoglochin* V.I.Krecz. | STA0027 | Molau & al. | 2329 | GB | Ecuador, Chimborazo | KJ513584 | KJ513492 | AY244520 |
| *Carex canescens* L. | - | Kaantonen | 156/94 | H | Finland | KP980061 | - | - |
| *Carex canescens* L. | - | Bond | s.n. | MTMG | Canada, Québec | - | - | AY757384 |
| *Carex capitata* Sol. | STA2653 | Starr & Thibeault | 6016 | CAN | United States | KJ513585 | KJ513493 | - |
| *Carex capitata* Sol. | STA1411 | Ford | 02379 | WIN | Canada | - | - | DQ115119 |
| *Carex dimorpholepis* Steud. | STA2645 | [not available] | MAK accession no. 99052601 | MAK | Japan, Nara Pref. | AB079435 | AB079422 | - |
| *Carex filicina* Nees | STA2459 | Ford & al. | 1247A | WIN | Vietnam, Lao Cai | KP273682 | KP273736 | KP273608 |
| *Carex hypolytroides* Ridl. | STA2442 | Ford & al. | 1255A | WIN | Vietnam, Lao Cai | KP273688 | KP273742 | KP273610 |
| *Carex kucyniakii* Raymond | STA2351 | Ford & al. | 1261A | WIN | Vietnam, Lao Cai | KP273693 | KP273747 | KP273615 |
| *Carex microglochin* Wahlenb. | STA2658 | Starr & al. | 10-008 (P5-9) | CAN | Argentina, Tierra del Fuego | KP273698 | KP273752 | - |
| *Carex microglochin* Wahlenb. | STA0106 | Starr & Scott | 98017 | FHO | Scotland, County of Perth | - | - | AY244518 |
| *Carex myosuroides* Vill. | B1518 | Jones | 146 | UBC | Canada, British Columbia | KJ513622 | KJ513529 | - |
| *Carex myosuroides* Vill. | STA0101 | Playford & al. | 9084 | FHO | France, Hautes-Alpes/Sa voie | - | - | AH012966 |
| *Carex myosurus* Nees | STA2456 | Ford & al. | 1246A | WIN | Vietnam, Lao Cai | KP273700 | KP273754 | KP273620 |
| *Carex nardina* (Hornem.) Fr. | - | Aiken & al. | 86-091 | CAN | Canada, Nunavut | FJ548120 | - | - |
| *Carex nardina* (Hornem.) Fr. | STA0839 | Ford & al. | 02230 | WIN | Canada, Manitoba | - | - | DQ115221 |
| *Carex phleoides* Cav. | - | Danton | G-(1377)-1142 | [not available] | [not available] | - | AM999972 | - |
| *Carex phleoides* Cav. | STA0034 | Vann | s.n. | FHO | Chile, Chiloé | - | - | AH010381 |
| *Carex plantaginea* Lam. | STA0178 | Waterway | 2000.002 | MTMG | Canada, Québec | - | - | AY757674 |
| *Carex pulicaris* L. | STA0105 | Starr & Scott | 98001 | FHO | England, North Yorkshire | KJ513590 | KJ513576 | AY242019 |
| *Carex siderosticta* Hance | STA0733 | Léveillé-Bourret | 545 | CAN | Garden | KJ513592 | KJ513499 | - |
| *Carex siderosticta* Hance | - | Waterway | 2004.268 | MTMG | MTMG | - | - | DQ998892 |
| *Carex speciosa* Kunth | STA2417 | Ford & al. | 1236A | WIN | Vietnam, Lao Cai | KP273706 | KP273760 | KP273625 |
| *Khaosokia caricoides* D.A.Simpson et al. | STA0387 B | Middleton & al. | 4071 | MICH | Thailand, Surat Thani | JX065087 | JX074643 | - |
| *Dulichium arundinaceum* | STA0154 | Ford & Punter | 94233 | FHO | Canada, Manitoba | JX065083 | JX074639 | - |
| Pers. var.<br>*arundinaceum* |  |  |  |  |  |  |  |  |
| *Dulichium*<br>*arundinaceum*<br>Pers. var.<br>*arundinaceum* | STA2469 | Bergeron<br>& al. | 81113 | CAN | Canada,<br>Québec | - | - | KP705281 |
| *Eleocharis*<br>*acicularis* (Willd.)<br>Schult. | d484 | Fields | 2583 | WIS | United<br>States,<br>Wisconsin | KJ513595 | KJ513502 | - |
| *Erioscirpus*<br>*comosus* (Wall.)<br>Palla | STA2092 | Hing & al. | 22413 | A | China,<br>Yunnan | KJ513619 | KJ513526 | - |
| *Erioscirpus*<br>*comosus* (Wall.)<br>Palla | - | Ikeda & al. | 4032 | TI | Nepal | - | - | KM462231 |
| *Eriophorum*<br>*angustifolium*<br>Honck. | STA1777 | Scoggan | 10947 | CAN | Canada | KJ513597 | KJ513504 |  |
| *Eriophorum*<br>*angustifolium*<br>Honck. | STA2547 | Keleher | 755 | CAN | Canada | - | - | KP705276 |
| *Eriophorum*<br>*vaginatum* subsp.<br>*spissum* (Fern.)<br>Hultén | STA1804 | Porsild | 12 | CAN | Canada | KJ513614 | KJ513521 | - |
| *Eriophorum*<br>*vaginatum* subsp.<br>*spissum* (Fern.)<br>Hultén | STA2607 | Léveillé-Bourret | 632 | DAO | Canada | - | - | KP705278 |
| *Eriophorum*<br>*virginicum* L. | STA1807 | Dickson &<br>Brunton | 3214 | CAN | Canada | KJ513617 | KJ513524 | - |
| *Eriophorum*<br>*virginicum* L. | STA2608 | Léveillé-Bourret | 633 | DAO | Canada | - | - | KP705269 |
| *Scirpus atrovirens*<br>Willd. | dS043 | Spalink | 186 | WIS | United<br>States,<br>Ohio | KJ513630 | KJ513537 | - |
| *Scirpus cyperinus*<br>(L.) Kunth | STA1777 | Lindsey | 1025 | CAN | Canada,<br>Ontario | JX065092 | JX074648 | - |
| *Scirpus pendulus*<br>Muhl. | STA1878 | Cruise | 1388 | CAN | Canada | KJ513649 | KJ513556 | - |
| *Scirpus pendulus*<br>Muhl. | STA2569 | Léveillé-Bourret | 611 | DAO | Canada | - | - | KP705270 |
| *Scirpus rosthornii*<br>Diels | STA2437 | Ford & al. | 1260A | WIN | Vietnam,<br>Lao Cai | newly<br>submitted | - | newly<br>submitted |
| *Trichophorum*<br>*alpinum* (L.) Pers. | STA1815 | Spetzman | 4941 | CAN | United<br>States,<br>Alaska | JX065093 | JX074649 | KP705266 |
| *Trichophorum*<br>*cespitosum* (L.)<br>Schur | STA1819 | Aiken &<br>Iles | 02-048 | CAN | Canada,<br>Nunavut | KJ513657 | KJ513564 | KP705267 |

### Online Appendix 1

Characteristics of loci used in anchored phylogenomics analyses after masking and trimming of alignments. AVE: average value for all loci, MIN: minimum value for all loci, MAX: maximum value for all loci, Locus#: locus identification number, New#: new number assigned to locus for this analysis (necessary because of paralogs), cov: average coverage across all species, ntax: number of taxa possessing the locus, flank%: percentage of total locus length estimated to be in the “flanking region”, aveC: average number of copies of locus across species, before orthology filtering, maxC: maximum number of distinct copies of locus, before orthology filtering, len: alignment length, var: number of variable characters, pi: number of parsimony informative characters, GC%: percentage of GC across all non-gapped bases, amb%: percentage of ambiguous bases, miss%: percentage of missing data.

### Online Appendix 2

Saturation plot as calculated in DAMBE, showing GTR distance estimated for the 1/3 fastest-evolving sites in relation to the number of transitions and transversions at these sites. Regression lines (forced through the origin) and slope coefficients are shown. Dashed line indicates slope of 1.

### Online Appendix 3

Single shortest tree found in PAUP* maximum parsimony searches. Support as MP jackknife, and ACCTRAN branch lengths.

### Online Appendix 4

Maximum likelihood tree in RAxML searches. Support as ML bootstrap.

